# An Acquisition-agnostic k-Space Denoising Method for High-dimensional Imaging

**DOI:** 10.64898/2026.01.20.699791

**Authors:** Ludwig Sichen Zhao, Manuel Taso, Jay A. Gottfried, John A. Detre, M. Dylan Tisdall

**Author notes:** **Correspondence** M. Dylan Tisdall.

## Abstract

**Purpose:** High-dimensional and dynamic MRI are often limited by thermal noise, particularly in accelerated acquisitions. Although image-domain low-rank denoising methods (e.g., MP-PCA and NORDIC) are effective, their reliance on stationary noise distributions limits applicability to non-Cartesian sampling and advanced reconstructions. This work introduces Magnetic Resonance *K*-space Local low-rank Estimation for Attenuating Noise (MR KLEAN), a k-space low-rank denoising framework agnostic to acquisition trajectory and reconstruction strategy.

**Theory and Methods:** MR KLEAN exploits local low-rank structure in multichannel, high-dimensional k-space. Data are prewhitened using a noise-only scan to enforce independent and identically distributed, zero-mean, unit-variance noise. Casorati matrices from local k-space patches are denoised by singular-value thresholding, with thresholds set via Monte-Carlo simulations under known noise statistics. MR KLEAN was evaluated in (1) a Cartesian 3D FLASH phantom study, (2) an ASL study with spiral readout and compressed sensing reconstruction to assess generalizability and preservation of temporal information via resting-state connectivity analysis, and (3) an accelerated cardiac cine study assessing performance under rapid temporal dynamics.

**Results:** MR KLEAN increased SNR and CNR in phantom studies. In vivo ASL showed reduced noise in perfusion images, improved relative SNR, and substantially enhanced resting-state networks detection. In cardiac imaging, noise was reduced and delineation of fine anatomical features improved while temporal fidelity was preserved across cardiac phases.

**Conclusion:** MR KLEAN provides robust, acquisition- and reconstruction-agnostic k-space denoising, improving image quality and allowing flexible spatial-temporal trade-offs. Results further support that high-dimensional k-space data retain intrinsic local low-rank structure analogous to image-space despite temporal signal variations.

## 1 INTRODUCTION

The inherent low signal-to-noise ratio (SNR) of MRI is exacerbated in high-dimensional or time-resolved (dynamic) imaging, where maintaining adequate SNR usually requires trading spatial resolution, temporal resolution, or additional encoding dimensions against overall acquisition time. For example, diffusion MRI (dMRI) and BOLD fMRI acquired with EPI are widely used to map brain structural and functional connectivity, respectively. Despite their different contrasts, these rapid imaging modalities both share a fundamental limitation of SNR^1,2,3,4^. Arterial spin labeling (ASL), which directly quantifies cerebral perfusion and serves as an alternative biomarker of brain function to BOLD fMRI, is similarly constrained by its inherent low SNR, even with repeated measurements^5,6,7,8,9^. This limitation is not confined to neuroimaging; other dynamic MRI applications throughout the body can suffer also exhibit low-SNR characteristics due to demands of high temporal resolution, motion robustness, and specific contrast mechanisms^10,11,12,13^.

Rather than addressing SNR via trading off resolution against scan time, an alternative is to suppress thermal noise through post-acquisition denoising. Denoising can be expressed as an explicit filtering operation applied to the data before or after reconstruction, but is also sometimes expressed implicitly as a regularization step within the image reconstruction algorithm. In this work, we explore an explicit denoising strategy, operating on raw k-space data and applied before any subsequent reconstruction steps. In particular, we propose a denoising strategy that models MRI data as low-rank matrices and attenuates thermal noise via singular value thresholding (SVT). Our choice of this denoising strategy takes inspiration from the Noise Reduction with Distribution Corrected (NORDIC) PCA method that has gained considerable attention for efficiently and effectively removing thermal noise from diffusion and BOLD fMRI data^3,14^. However, we note a key limitation of NORDIC: it operates on local patches of reconstructed image-space data, prewhitening the coil-combined images using a *g*-factor map before applying SVT. This requirement for a *g*-factor map makes it challenging to apply in combination with non-linear and iterative reconstruction techniques such as compressed sensing or deep learning. While several variants have been proposed that perform denoising prior to reconstruction to avoid the need for a *g*-factor map, they are limited to specific Cartesian trajectories^15,16^.

To address this shortcoming of the NORDIC method more generally, we propose to apply the SVT step directly in k-space, before image reconstruction. By operating in k-space, where thermal noise is well-characterized without the need for *g*-factor maps, the efficiency benefits of the patch-wise SVT denoising strategy can be generalized to a wider range of sequences, and also take advantage of the additional redundancy inherent in multi-channel data. We call this strategy, Magnetic Resonance *K* -space Local low-rank Estimation for Attenuating Noise (MR KLEAN).

We evaluated MR KLEAN across different anatomies and k-space trajectories, including those where directed g-factor estimates are not easily available, thus limiting the applicability of NORDIC. MR KLEAN proved to be a robust and generalizable denoising technique for multichannel and high-dimensional MRI. Moreover, the results provide empirical evidence supporting the presence of low-rank structure in Casorati matrices constructed from k-space patches. By leveraging this property, MR KLEAN improves SNR without sacrificing spatial or temporal resolution, making it a broadly applicable and effective denoising solution for highdimensional, low-SNR MRI that is agnostic to both acquisition and reconstruction methods.

## 2 THEORY

### 2.1 Singular Value Thresholding (SVT) for Denoising

The key intuition of SVT denoising is that structured matrices can be constructed from the MRI data that can then be effectively compressed in a low-rank representation, separating the true MRI signal from noise. A wide variety of strategies have been proposed for constructing matrices with appropriate structure to support SVT denoising. Much of the prior work applying SVT in MRI has focused on imposing global or local low-rank (LLR) structure on image-domain data^17,18,19,3,14^. Moreover, when expressed as a regularization term, image-domain LLR methods have also been applied with complementary priors, most commonly sparsity, within the reconstruction objective to improve robustness and enable accelerated dynamic imaging under a range of undersampling strategies^20,21,22,23,24^.

While the methods described above apply the low-rank constraint in image-space, others have observed that k-space also has low-rank structure. Partially separable function models formulate reconstruction as recovering a global low-rank Casorati representation directly in k-space^25,26^. A related method, LORAKS, imposes LLR constraints in k-space based on specific image-space features (e.g., finite support and smooth phase) by constructing Hankel matrices from patches of k-space^27^. Similar structures have also been presented as regularization terms in reconstructing undersampled multi-channel data^28,29^, and additionally in combination with further sparsifying transforms^30^.

In this work, we focus on applying SVT denoising to high-dimensional MRI data – such as multiecho acquisitions indexed by TE, dMRI indexed by *q*-space sampling, or fMRI indexed by time – where redundancy across time, contrast, etc. can be exploited in addition to spatial structure. As noted above, we take inspiration from the NORDIC algorithm, which creates patch-wise Casorati matrices of image/time-domain data after reconstruction and pre-whitening and then applies SVT to denoise them. In MR KLEAN, we instead create patch-wise Casorati matrices of k-space data and apply SVT to these before reconstruction. This specific choice offers two unique benefits: first, unlike NORDIC it does not require estimation of spatially varying noise statistics, as the thermal noise is spatially stationary in k-space; second, the application of SVT as a denoising method fully separated from any regularization terms applied in reconstruction allows the SVT denoising to be easily generalized to a wide variety of existing pulse sequences.

### 2.2 Proposed Denoising Method

An overview of the proposed MR KLEAN processing pipeline is shown in Figure 1; the individual steps are described in the following subsections.

**FIGURE 1:**
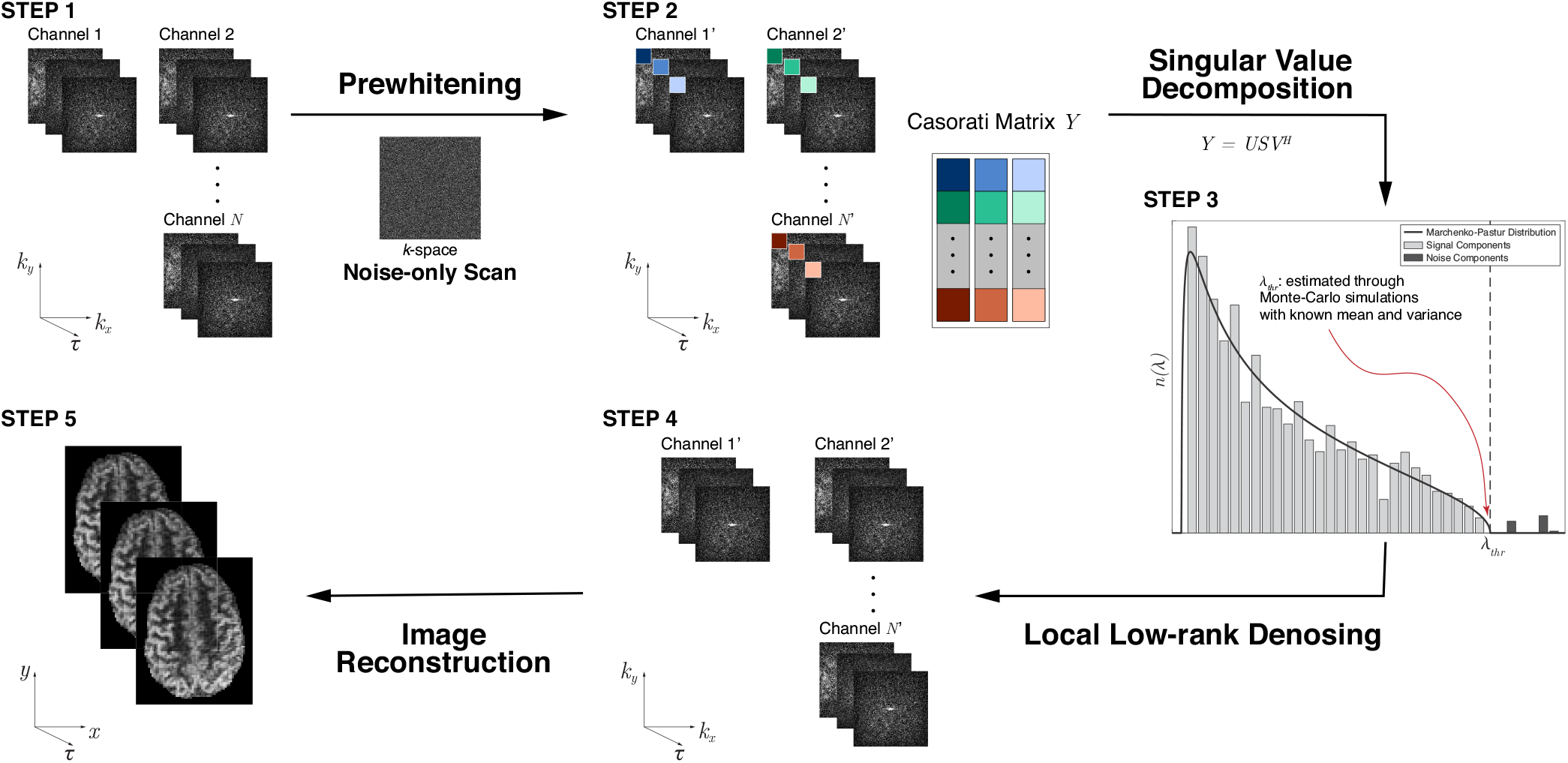
Flowchart of the proposed MR KLEAN k-space denoising pipeline. Multichannel k-space data is prewhitened using a noise-only scan (Section 2.2.1), concatenated into Casorati matrices, and denoised via LLR (Section 2.2.2) with a known threshold (Section 2.2.3) to separate signal and noise components prior to image reconstruction.

#### 2.2.1 Prewhitening

In a single channel of k-space data, the noise is well-described as additive, circular complex Gaussian, i.i.d., and zero-mean, thus satisfying the assumptions of random matrix theory. When using phased-array coils, although the noise in k-space is generally correlated across channels, the noise can be decorrelated through prewhitening and therefore meet i.i.d. and zero-mean conditions.

To estimate the noise covariance, a noise scan 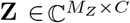 is acquired, where *M*_*Z*_ is the number of noise samples collected from an individual channel and *C* is the number of channels. The prewhitened noise data **Z**_*w*_ = **ZW** shall satisfy the following condition:

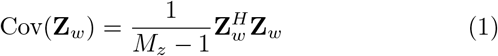

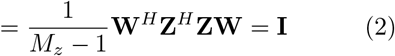

Therefore, **W** can be computed through

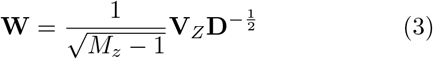

where 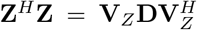. **D** is the diagonal matrix of eigenvalues of **Z**^*H*^ **Z** and **V**_*Z*_ is the unitary matrix whose columns are the associated eigenvectors. Applying the transform ***s***^*′*^ = **Z*s***, where an acquired k-space sample is the *C*-vector ***s*** and it’s prewhitened form is ***s***^*′*^, produces prewhitened measurements.

#### 2.2.2 LLR Denoising

Let the prewhitened complex k-space data from channel *j* ∈ {1, …, *C*} be denoted by 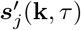. **k** is the location in 3D k-space, *τ* ∈ {1, …, *N*} indexes the “time” dimension (i.e., TE, volume, diffusion weighting), and *C* is the number of channels. k-space data at each “time” *τ*, within each channel *j*, is partitioned into non-overlapping patches of size *p*_1_ × *p*_2_ × *p*_3_. Each patch, indexed with *l*, is vectorized to form **y**_*τ,j,l*_. These vectors are then assembled into a Casorati matrix **Y**_*l*_ ∈ ℂ^*M ×N*^, as follows:

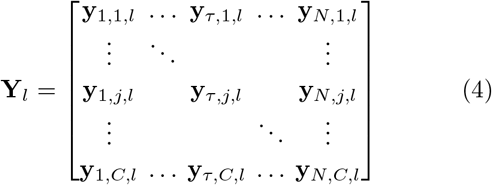

where *M* = *p*_1_*p*_2_*p*_3_ · *C*.

We first perform singular value decomposition (SVD) on **Y**_*l*_:

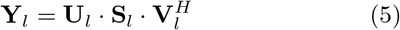

To denoise the data, we select a threshold 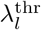 to generate 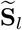. The denoised Casorati matrix is then obtained through

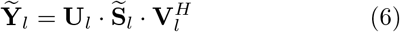

To compute the denoised k-space data, we reassemble all patches into their original matrix form and apply the inverse prewhitening transform **W**^−1^.

#### 2.2.3 Threshold Estimation

Across the various approachs to SVT denoising that have been previously preseted in the context of MRI, selecting the singular value cutoff (or effective rank) remains a recurring practical challenge and is often treated as a tunable hyperparameter whose value varies across applications and acquisition settings. While some work has developed principled, model-based rules for tuning singular-value shrinkage under explicit noise assumptions^31^, a more commonly used alternative is Marchenko–Pastur Principal Component Analysis (MP-PCA), which integrates LLR modeling with random matrix theory to directly estimate the singular value cutoff given^32,33^. However, it assumes that noise is (1) independent and identically distributed (i.i.d.) and (2) zero-mean. These conditions are often violated in magnitude MR images, particularly when using phased-array coils or parallel imaging reconstruction. In NORDIC, this issue is addressed via the g-factor map, allowing point-wise estimation of noise statistics. In MR KLEAN, the process can be simplified due to the stationary noise-statistics of k-space.

Specifically, for each all patches *l* with a shared patch size, a single shared threshold 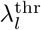 is estimated through Monte-Carlo simulations. We generate a set of random matrices – each having the same dimensions as **Y**_*l*_ – with entries drawn i.i.d. from a zero mean and unit variance Gaussian distribution. We then compute the largest singular value, *σ*_max_, of each randomly generated matrix, and define the threshold as the sample mean of these largest singular values, 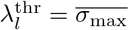.

## 3 METHODS

### 3.1 Participants

Two healthy participants provided informed consent under a protocol approved by the University of Pennsylvania Institutional Review Board (IRB #817383): one female (32 years old) for the ASL study and one male (45 years old) for the cardiac study.

### 3.2 Phantom Study

We first conducted a phantom study to evaluate the denoising performance of MR KLEAN under stationary conditions using a standard Cartesian readout, isolating the effects of thermal noise without contributions from physiological or motion-related noise sources.

An ACR MRI phantom (JM Specialty Parts, San Diego, CA, USA) was scanned to evaluate MR KLEAN performance under controlled conditions. A 3D FLASH sequence (TR/TE = 6.6/3.3 ms, flip angle = 15 deg, field of view [FOV] = 200 x 200 mm^2^, 1.04 mm isotropic resolution, 35 volumes) was acquired on a 3 T MRI scanner (MAGNETOM Prisma, Siemens Healthineers, Erlangen, Germany) using the vendor-provided 20-channel head and neck coil^34^. To simulate varying noise levels, the scan was acquired twice, once with a bandwidth of 360 Hz/pixel and a second time at 1630 Hz/pixel. Three volumes of a noise-only scan were acquired with identical parameters but with the RF pulses disabled.

To quantify denoising performance, we measured SNR and contrast-to-noise ratio (CNR) as a function of the number of averaged volumes for both low and high readout bandwidths, as defined in Section 3.7.1.

We also examined the effect of the number of coil channels on denoising performance: from the same dataset, images were first reconstructed with only a subset of coil channels most relevant to the region of interests (ROIs), after which additional channels were gradually incorporated. Denoising performance was evaluated using relative SNR and relative CNR, defined as the ratio of the metric after to before denoising, computed across all volumes (Section 3.7.1).

### 3.3 ASL Study

To demonstrate the generalizability of MR KLEAN, we applied it to ASL data acquired and reconstructed using a non-Cartesian spiral trajectory and compressed-sensing reconstruction. This dataset reflects dynamic brain imaging with slow temporal variations (in this case, cerebral perfusion changes due to resting-state brain activity). We evaluated denoising performance and whether temporal information was preserved by examining both averaged perfusion images and temporal signal characteristics.

ASL data were acquired on a 3 T scanner (MAGNETOM Prisma, Siemens Healthineers, Erlangen, Germany) using the vendor-provided 64-channel head and neck coil with an unbalanced pseudocontinuous ASL (pCASL)^35,36^ combined with background-suppressed, 3D accelerated Stack-of-Spirals (SoS) turbo spin echo (TSE) readout^37^ (TR/TE = 5000/9.4 ms, labeling duration = 3 s, post-labeling delay = 1.2 s, flip angle = 22.5 deg, *G*_*max*_*/G*_*avg*_ = 3.5/0.5 mT/m, 97.5% background suppression, FOV = 220 × 220 mm^2^, 2 mm isotropic resolution, 2 shots, 15 label–control pairs). Two volumes of a noise-only scan were also acquired with identical parameters but with the RF pulse disabled.

Fast 3D FLASH scans with matching FOV and resolution (TR/TE = 5/2 ms, bandwidth = 1180 Hz/pixel) were acquired for coil sensitivity estimation^34^.

### 3.4 Cardiac Study

We applied MR KLEAN to an accelerated cine cardiac acquisition to evaluate its denoising performance under conditions of rapid temporal variation.

Cine cardiac MRI was performed on a 1.5 T scanner (MAGNETOM Avanto^*fit*^, Siemens Healthineers, Erlangen, Germany) using the vendor-provided 18-channel body coil in combination with a 32-channel spine coil^38^. Retrospectively ECG-gated, breath-held acquisitions were obtained in the short-axis orientation using a 2D segmented balanced steady-state free precession (bSSFP) sequence (TR/TE = 34.65/1.26 ms, flip angle = 65 deg, FOV = 360 × 270 mm^2^, matrix = 240 × 204, slice thickness = 7 mm, 12 slices, bandwidth = 947 Hz/pixel, in-plane GRAPPA acceleration factor = 4 with 44 reference lines). A noise-only scan was acquired using identical parameters, except that only six slices were collected and the RF pulses were disabled.

### 3.5 Denoising

MR KLEAN was applied prior to any image reconstruction, following the algorithm described in Section 2.2.

For the 3D FLASH phantom study, the patch size was set to 6 voxels (*k*_*x*_) × 6 voxels (*k*_*y*_) × 6 voxels (*k*_*z*_). For the cardiac study, a patch size of 4 voxels (*k*_*x*_) × 4 voxels (*k*_*y*_) × 1 voxel (*k*_*z*_) was used.

For the ASL study, which used a spiral acquisition, the construction of **y**_*τ*_ was modified to account for the non-Cartesian sampling pattern. A patch was defined on each spiral arm, consisting of up to 40 consecutive acquired readout points (or fewer if fewer than 40 points remained on the given spiral) across up to 10 consecutive *k*_*z*_ partitions (or fewer if fewer than 10 slices were available). For each coil, **y**_*τ*_ was vectorized first along the spiral readout and then along *k*_*z*_. MR KLEAN was then applied patch by patch on each individual spiral. All label and control images were denoised within the same time series.

### 3.6 Image Reconstruction

All images were reconstructed offline from raw k-space data using MATLAB (R2020b, MathWorks Inc., Natick, MA, USA). For both the 3D FLASH phantom study and the cine cardiac study, coil channels were sum-of-squares combined. For ASL data, coil sensitivities were first estimated from the 3D FLASH data using ESPIRiT (*σ* = 0.9, *c* = 0.01)^39^. The images were then reconstructed using a LLR regularized compressed sensing (CS) approach^40^ (*λ* = 0.001, 100 iterations) implemented in the BART toolbox (0.9.00)^41,42^. The reconstructed ASL volumes were subsequently motion-corrected using FSL (6.0.6.2, FMRIB Software Library, University of Oxford, Oxford, UK)^43^. Perfusion-weighted images were then generated by subtracting the label volume from the corresponding control volume.

### 3.7 Data Analysis

#### 3.7.1 SNR, Relative SNR, CNR and Relative CNR

For each acquired 4D image series ***m***(**r**, *τ*), as defined in Section 2.1, we first averaged across the volumes (indexed by *τ*) to obtain a single mean 3D image for subsequent analysis.

SNR was computed within an ROI as the ratio of the mean signal intensity to the standard deviation of the signal across voxels in the ROI:

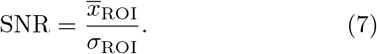

Relative SNR was defined as the ratio of SNR values between two image series computed within the same ROI.

CNR was computed as

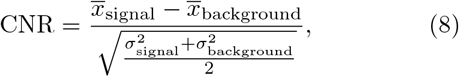

where 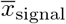 and 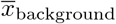 denote the mean intensities within the predefined signal and background regions, respectively, and 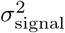 and 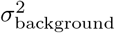 are the corre-sponding variances across voxels within those regions.

Relative CNR was defined as the ratio of CNR values between two image series computed using the same signal and noise regions.

#### 3.7.2 Seed-Based Connectivity (SBC) Maps

For the ASL study, SBC maps were computed by first extracting the mean time series from the perfusion images within the ROI. The voxel-wise Pearson correlation coefficient (*r*) between the ROI time series and the time series of each voxel was then calculated across all volumes to generate the connectivity map.

The seed region was the right posterior cingulate cortex (PCC). Accordingly, the ROI was defined as a 6-mm-radius sphere centered at (0, -53, 26) in Montreal Neurological Institute (MNI) space, consistent with prior work^44,45^. The MNI template (ICBM 152 Nonlinear Asymmetrical, version 2009c; MNI-ICBM2009c)^46,47^ was registered to the subject’s ASL proton density (*M*_0_) image^48^ using Advanced Normalization Tools (ANTs, 2.3.5)^49,50^ to estimate the spatial transformation, which was then applied to warp the ROI from MNI space into subject space.

## 4 RESULTS

### 4.1 Phantom Study

Figure 2A demonstrates a representative volume before and after denoising. Visually, MR KLEAN reduces background grain and more clearly delineates fine structural features. Quantitative results are shown in Figures 2B and 2C: as expected, increasing the readout bandwidth led to noisier images, whereas SNR and CNR increased with the number of volumes averaged and eventually approached a plateau.

**FIGURE 2:**
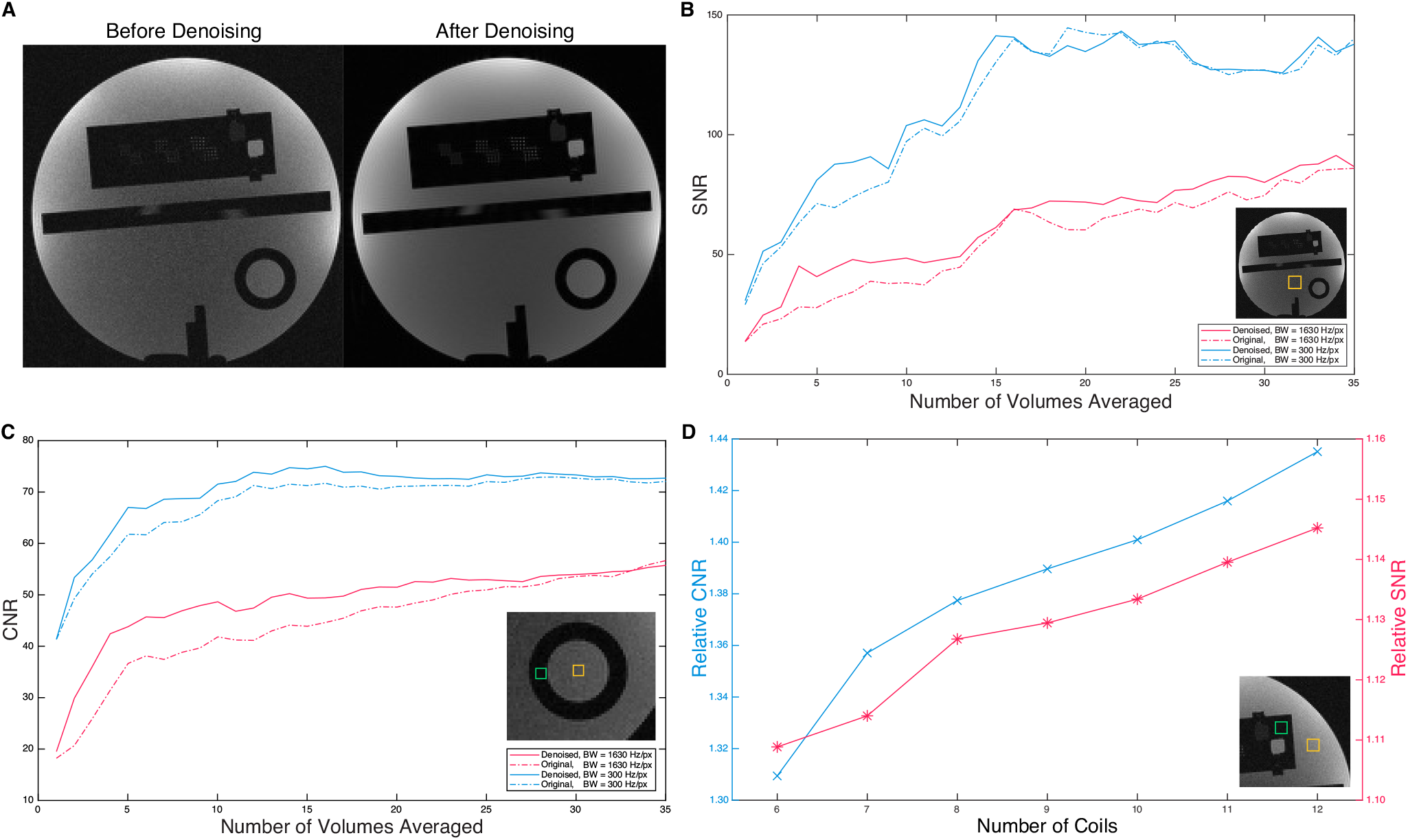
Comparison of images before and after denoising in the phantom study. A) A representative slice from a single volume of the ACR phantom before and after applying MR KLEAN across all 35 volumes. B) SNR before (dashed lines) and after (solid lines) applying MR KLEAN for different numbers of averaged volumes at low readout bandwidth (300 Hz/pixel, blue) and high readout bandwidth (1630 Hz/pixel, red). The ROI used for SNR measurement is indicated by the yellow square in the inset. C) CNR before (dashed lines) and after (solid lines) applying MR KLEAN for different numbers of averaged volumes at low readout bandwidth (300 Hz/pixel, blue) and high readout bandwidth (1630 Hz/pixel, red). The signal and background ROIs used for CNR estimation are shown as yellow and green squares, respectively, in the inset. D) Relative SNR (ratio of SNR after to before applying MR KLEAN to all volumes, red) and relative CNR (ratio of CNR after to before applying MR KLEAN to all volumes, blue) for the high readout bandwidth (1630 Hz/pixel) across different numbers of coil channels. The inset shows the ROIs used for measurements: the yellow square denotes the ROI for SNR measurement and the signal region for CNR estimation, while the green square indicates the background region used for CNR estimation.

For single-volume acquisitions, MR KLEAN provided minimal improvement in SNR at either bandwidth.

In contrast, with two or more volumes, MR KLEAN produced substantial SNR gains, particularly at the higher readout bandwidth. With additional volumes being averaged, the SNR improvement increased further, plateauing at approximately 18 volumes for the low-bandwidth condition, while continuing to increase for the high-bandwidth condition.

CNR followed a similar trend. Overall, the low-bandwidth acquisition yielded higher CNR, and MR KLEAN further increased CNR as the number of averaged volumes increased. Although improvements were minimal for single-volume data, substantial gains were observed for multivolume acquisitions, demonstrating effective suppression of thermal noise. Notably, for the low bandwidth condition, the CNR measured after denoising with 8 volumes was approximately comparable to that obtained with 20 volumes before denoising, suggesting that MR KLEAN could effectively reduce the required acquisition time by more than half.

Finally, Figure 2D shows that both relative SNR and relative CNR increased consistently with the number of coil channels. Even when restricted to the single channel most relevant to the ROIs, MR KLEAN remained effective, yielding a relative CNR of 1.64 and a relative SNR of 1.45.

### 4.2 ASL Study

Figure 3A shows representative perfusion images before and after denoising across varying numbers of averaged label–control pairs, while Figure 3B presents the corresponding quantitative analysis using relative SNR for both cortical and white matter regions. Similar to the phantom study, a single-volume perfusion image provided a limited opportunity for denoising. However, with two or more volumes, MR KLEAN achieved noticeable noise suppression, resulting in an improved perfusion image and SNR.

**FIGURE 3:**
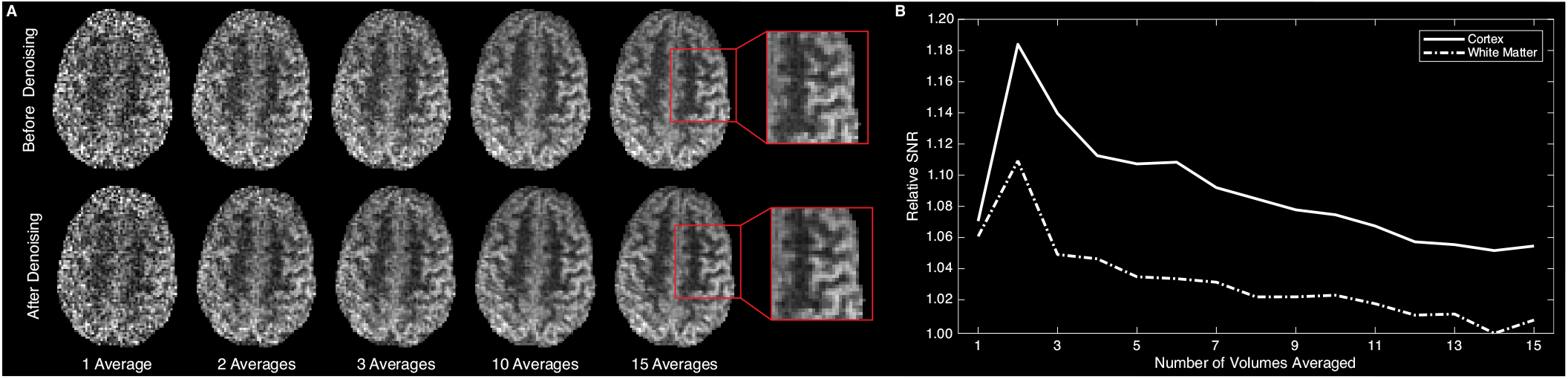
Comparison of Images before and after denoising in the ASL study. A) A representative slice of perfusion-weighted images averaged over different numbers of label–control pairs, shown before (top row) and after (bottom row) MR KLEAN denoising. B) Relative SNR (ratio of SNR after to before applying MR KLEAN) across different numbers of averaged label–control pairs for cortex and white matter regions in the representative slice.

To evaluate the impact of MR KLEAN on temporal dynamics, we next examined functional connectivity of the resting-state by seeding the right PCC and computing voxel-wise correlation maps before and after denoising (Figure 4). Without denoising, connectivity was weak and spatially confined. In contrast, after applying MR KLEAN, clear network patterns emerged that closely resembled the default mode network (DMN), indicating that MR KLEAN not only preserved temporal information but also improved sensitivity to physiologically significant fluctuations in brain activity.

**FIGURE 4:**
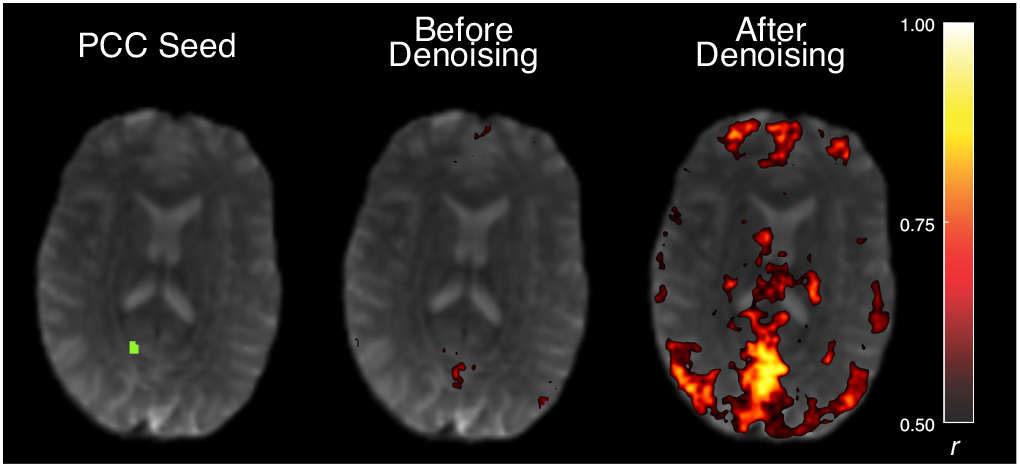
Seed-based connectivity before and after denoising. The seed region is shown in green in the right posterior cingulate cortex.

These results show that MR KLEAN effectively improves both the spatial quality and the sensitivity of the temporal dynamics of ASL data acquired with spiral readouts, demonstrating a generalizability to non-Cartesian, dynamic imaging applications.

### 4.3 Cardiac Study

Figure 5 shows three consecutive short-axis slices from a cine cardiac scan before and after denoising, demonstrating that MR KLEAN improves image quality under rapid temporal variation. Visually, MR KLEAN substantially reduced noise, particularly in the myocardium, while preserving anatomical detail and enhancing the definition of epicardium (red arrows), papillary muscles (blue arrows) and ventricular trabeculae (yellow arrows). A motion clip demonstrating the same three slices across all 23 cardiac phases is provided in Supporting Information Video S1, showing that MR KLEAN effectively removes noise from individual volumes while preserving the temporal fidelity of cardiac motion.

**FIGURE 5:**
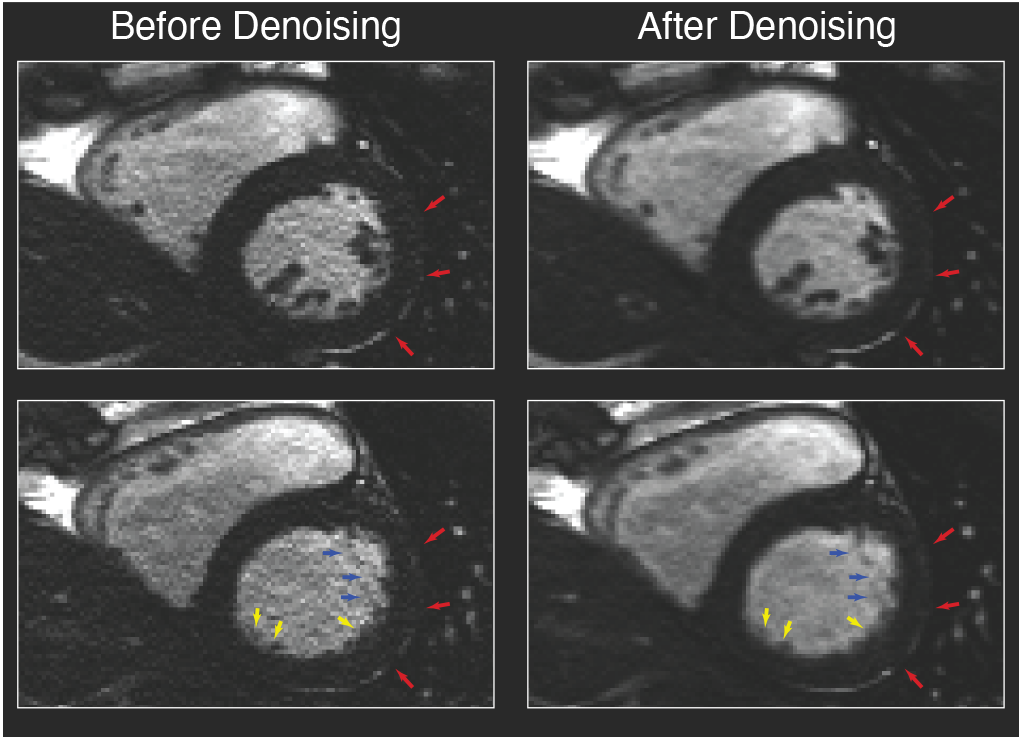
Comparison of images before and after denoising in the cardiac study. Three consecutive short-axis slices from a single cine volume are shown before and after applying MR KLEAN. Pre-scan normalization was applied for visualization purposes. Red arrows: epicardium. Blue arrows: papillary muscle. Yellow arrows: ventricular trabeculae.

## 5 DISCUSSION

In this work, we have describe and evaluated the use of an SVT denoising filter exploiting LLR structure in k-space. This particular formulation of SVT denoising can be viewed as a close relative of the NORDIC denoising algorithm, but in our case applied to k-space instead of image-space. Specifically, we have dmonstrated that multivolume k-space data also exhibits a LLR structure when structured as non-overlapping patches in the form of a Casorati matrix and that thermal noise can be effectively removed by thresholding singular values according to known noise statistics. The MR KLEAN algorithm has several key advantages that enable it to be both efficient and generally applied to many MRI sequences. First, by operating in k-space, before any image reconstruction, the noise statistics are well-known and stationary, enabling the threshold to be efficiently estimated based only on the patch dimensions. Second, by using nonoverlapping patches in a Casorati matrix structure, MR KLEAN only performs one denoising operation for each point in k-space, compared to other structured matrix constructions that rely on overlapping patches to exploit data redundancy. Third, by expressing the denoising step as a separate operation, instead of as a regularization term in the reconstruction, MR KLEAN can denoise data regardless of the subsequent reconstruction algorithm employed.

We have demonstrated the performance of MR KLEAN through three experiments designed to evaluate its effectiveness and robustness under progressively more complex imaging conditions. First, as shown in Figure 2, MR KLEAN effectively removed thermal noise and improved image quality in a controlled phantom study, free from physiological noise or motion effects. Next, an ASL study (Figures 3 and 4) demonstrated two key advantages of MR KLEAN: (1) its generalizability across non-linear and iterative reconstruction methods, and (2) its ability to preserve and improve temporal dynamics in data with relatively slow-moving spatial structures (i.e., neuroimaging in a compliant participant). Lastly, we extended our evaluation to dynamic imaging scenarios using an accelerated bSSFP cine cardiac scan (Figure 5). Despite rapid temporal changes due to cardiac motion, MR KLEAN effectively suppressed noise within each time frame while maintaining temporal fidelity, demonstrating that the automatically chosen denoising threshold does not introduce blurring across the temporal dimension. Collectively, these results highlight the versatility and robustness of MR KLEAN across acquisition strategies, readout trajectories, and temporal dynamics. Moreover, these findings provide strong evidence supporting our hypothesis that multivolume k-space data, regardless of the temporal changes, exhibit a LLR structure analogous to that observed in the image-space when structured as patchwise Casorati matrices.

When examining the effect of the number of volumes, we observed consistent trends in Figures 2B and 2C. For a single-volume acquisition, the SNR and CNR are identical before and after denoising. However, as more volumes are incorporated, the denoised images rapidly exceed the original SNR and CNR, with the improvement gradually approaching the levels obtained through conventional averaging. A similar trend can be found in Figure 3C, where the relative SNR equals 1 for a single volume, increases sharply with the addition of only a few volumes, and eventually converges to the value achieved through averaging alone.

These results can be explained by considering (4). When the number of volumes is limited, the Casorati matrix **Y**_*l*_ ∈ ℂ^*m×n*^ becomes tall (*m* ≫ *n*), degrading the separation between signal and noise singular values. In the extreme case where *n* = 1, **Y**_*l*_ contains only a single singular value, making it impossible for denoising to isolate a noise-only component; consequently, no meaningful denoising can be achieved. As additional volumes are incorporated, the separation between signal and noise improves, more ranks can be retained, and MR KLEAN becomes increasingly effective.

Finally, we examined the effect of the number of coil channels on MR KLEAN’s denoising performance (Figure 2D). MR KLEAN operates effectively on single-channel data, but incorporating additional coil channels further improves denoising performance. This trend is consistent with the intuition that multichannel acquisitions increase redundancy and strengthen lowdimensional structure in the data, thereby improving the effectiveness of LLR modeling. More broadly, these results suggest that MR KLEAN is most advantageous when additional dimensions (coils, time, repetitions) increase the intrinsic low-rank structure, even when the signal varies rapidly across those dimensions.

MR KLEAN represents a set of algorithm design choices within the broader family of low-rank MRI methods: operating in k-space instead of image-space; applying LLR instead of global low-rank constraints; constructing Casorati matrices from non-overlapping patches instead of other Hankell matrices or structures to express redundancy; estimating the threshold directly from matrix noise statistics instead of iteratively based on the measured data in each patch; and finally implementing the low-rank constraint as a separate filter before reconstruction rather than as a regularization constraint within the reconstruction.

While we eavaluted MR KLEAN using highdimensional (4D) imaging datasets in this study, its framework is equally applicable to three-dimensional acquisitions if acquired with multiple repetitions. For example, 3D structural or quantitative imaging protocols that rely on averaging – such as repeated short *T*_1_-weighted scans shown by Elliott et al. to improve precision in brain morphometry through repetition averaging – could also benefit from MR KLEAN denoising applied across repeated volumes^51^.

While MR KLEAN has demonstrated efficient and versatile denoising performance, several questions remain open for future investigation. As with other LLR-based methods, the selection of the optimal patch size remains heuristic and warrants systematic evaluation^52,53,54^. In this study, the Casorati matrix for Cartesian acquisitions was constructed following the conventional approach used in MP-PCA and NORDIC. For non-Cartesian acquisitions, such as the spiral readout in the ASL study, the patch construction was adapted to account for the sampling geometry. Nevertheless, additional optimization and validation of the non-Cartesian patch design are needed, despite the strong empirical performance observed in this work.

## 6 CONCLUSIONS

MR KLEAN is a low-rank denoising method that operates directly in k-space, providing an efficient, generalized denoising strategy for MRI data where the image-space algorithms are difficult to apply due to its preconditions. Taking advantage of the inherent statistical structure of high-dimensional and multichannel MRI data, MR KLEAN consistently suppresses thermal noise across diverse acquisition schemes while preserving spatial detail and temporal fidelity. In this study, we demonstrated its effectiveness in phantom, ASL, and cardiac imaging, further highlighting its robustness in varying contrast mechanisms, sampling trajectories, reconstruction strategies, and temporal dynamics. Collectively, these results establish MR KLEAN as a robust, acquisition- and reconstruction-agnostic denoising method that is broadly and easily applicable to highdimensional MRI acquisitions and may help mitigate image-quality degradation in accelerated imaging.

## ACKNOWLEDGMENTS

This work was supported by the National Institutes of Health under awards R01EB031080, R01AG080734, and R01DC018075. We thank the Center for Advanced Magnetic Resonance Imaging and Spectroscopy, RRID: SCR 022398, for their assistance with the MRI portion of the research. MT was employed by Siemens Medical Solutions USA during part of this work.

## Data Availability Statement

The source code and scripts used in this work are publicly available at https://github.com/luxwig/MR-CLEAN and archived at DOI:10.5281/zenodo.18311381. The software revision used for the results reported in this manuscript is SHA-1: 5f8bc99a9eaf8bf11ebbbe9ff8c368119f0c33a7.

## SUPPORTING INFORMATION

The following supporting information is available as part of the online article:

## Supporting Information Video S1

Video of cine images obtained before and after denoising. Two representative cardiac cine slices are shown before (left) and after (right) denoising.

## REFERENCES

1. Jones Derek K. Challenges and limitations of quantifying brain connectivity in vivo with diffusion MRI. Imaging in Medicine. 2010;2(3):341–355.

2. Manzano Patron Jose Pedro, Moeller Steen, Andersson Jesper L.R., Ugurbil Kamil, Yacoub Essa, Sotiropoulos Stamatios N.. Denoising diffusion MRI: Considerations and implications for analysis. Imaging Neuroscience. 2024;2:imag–2–00060.

3. Vizioli Luca, Moeller Steen, Dowdle Logan, et al. Lowering the thermal noise barrier in functional brain mapping with magnetic resonance imaging. Nature Communications. 2021;12(1):5181.

4. Triantafyllou Christina, Polimeni Jonathan R., Wald Lawrence L.. Physiological noise and signal-to-noise ratio in fMRI with multi-channel array coils. NeuroImage. 2011;55(2):597–606.

5. Hernandez-Garcia Luis, Aramendía-Vidaurreta Verónica, Bolar Divya S., et al. Recent Technical Developments in ASL: A Review of the State of the Art. Magnetic Resonance in Medicine. 2022;88(5):2021–2042.

6. Alsop David C., Detre John A., Golay Xavier, et al. Recommended implementation of arterial spin-labeled perfusion MRI for clinical applications: A consensus of the ISMRM perfusion study group and the European consortium for ASL in dementia. Magnetic Resonance in Medicine. 2015;73(1):102–116.

7. Garcia Dairon M., Duhamel Guillaume, Alsop David C.. Efficiency of inversion pulses for background suppressed arterial spin labeling. Magnetic Resonance in Medicine. 2005;54(2):366–372.

8. Detre John A., Wang Jiongjiong. Technical aspects and utility of fMRI using BOLD and ASL. Clinical Neurophysiology. 2002;113(5):621–634.

9. Detre John A., Leigh John S., Williams Donald S., Koretsky Alan P.. Perfusion imaging. Magnetic Resonance in Medicine. 1992;23(1):37–45.

10. Raptis Constantine A., Ludwig Daniel R., Hammer Mark M., et al. Building blocks for thoracic MRI: Challenges, sequences, and protocol design. Journal of Magnetic Resonance Imaging. 2019;50(3):682–701.

11. Wieben Oliver, Francois Christopher, Reeder Scott B.. Cardiac MRI of ischemic heart disease at 3 T: Potential and challenges. European Journal of Radiology. 2008;65(1):15–28.

12. Rajiah Prabhakar Shantha, François Christopher J., Leiner Tim. Cardiac MRI: State of the Art. Radiology. 2023;307(3):e223008.

13. Wen H., Denison T. J., Singerman R. W., Balaban R. S.. The Intrinsic Signal-to-Noise Ratio in Human Cardiac Imaging at 1.5, 3, and 4 T. Journal of Magnetic Resonance. 1997;125(1):65–71.

14. Moeller Steen, Pisharady Pramod Kumar, Ramanna Sudhir, et al. NOise reduction with DIstribution Corrected (NORDIC) PCA in dMRI with complex-valued parameter-free locally low-rank processing. NeuroImage. 2021;226:117539.

15. Demirel Omer, Moeller Steen, Vizioli Luca, et al. HighQuality 0.5mm Isotropic Functional MRI Using a Synergistic Combination of NORDIC Denoising and Deep Learning Reconstruction. In: Proceedings of the Joint Annual Meeting ISMRM-ESMRMB & ISMRT Annual Meeting, International Society for Magnetic Resonance in Medicine; 2022:3950; London, England, UK.

16. Moeller Steen, Olman Cheryl, Vizioli Luca, et al. NORDIC denoising before image reconstruction. In: Proceedings of the 2021 ISMRM & SMRT Annual Meeting & Exhibition, International Society for Magnetic Resonance in Medicine; 2021.

17. Trzasko J, Manduca A. Local versus Global Low-Rank Promotion in Dynamic MRI Series Reconstruction. ;.

18. Trzasko Joshua D.. Exploiting local low-rank structure in higher-dimensional MRI applications. In: Wavelets and Sparsity XV, SPIE; 2013:551–558.

19. Xu Lin, Wang Changqing, Chen Wufan, Liu Xiaoyun. Denoising Multi-Channel Images in Parallel MRI by Low Rank Matrix Decomposition. IEEE Transactions on Applied Superconductivity. 2014;24(5):1–5.

20. Zhao Bo, Haldar Justin P., Christodoulou Anthony G., Liang Zhi-Pei. Image reconstruction from highly under-sampled (k, t)-space data with joint partial separability and sparsity constraints. IEEE Transactions on Medical Imaging. 2012;31(9):1809–1820.

21. Lingala Sajan Goud, Hu Yue, DiBella Edward, Jacob Mathews. Accelerated Dynamic MRI Exploiting Sparsity and Low-Rank Structure: k-t SLR. IEEE Transactions on Medical Imaging. 2011;30(5):1042–1054.

22. Otazo Ricardo, Candés Emmanuel, Sodickson Daniel K.. Low-rank plus sparse matrix decomposition for accelerated dynamic MRI with separation of background and dynamic components. Magnetic Resonance in Medicine. 2015;73(3):1125–1136.

23. Ong Frank, Lustig Michael. Beyond Low Rank + Sparse: Multiscale Low Rank Matrix Decomposition. IEEE Journal of Selected Topics in Signal Processing. 2016;10(4):672–687.

24. Meyer Nolan K., Kang Daehun, Ahmed Zaki, et al. Locally Low-Rank Denoising of Multi-Echo Functional MRI Data With Application in Resting-State Analysis. Topics in Magnetic Resonance Imaging. 2023;32(5):37.

25. Liang Zhi-pei. SPATIOTEMPORAL IMAGINGWITH PARTIALLY SEPARABLE FUNCTIONS. In: 2007 4th IEEE International Symposium on Biomedical Imaging: From Nano to Macro, ; 2007:988–991.

26. Haldar Justin P., Liang Zhi-Pei. Spatiotemporal imaging with partially separable functions: A matrix recovery approach. In: 2010 IEEE International Symposium on Biomedical Imaging: From Nano to Macro, ; 2010:716– 719.

27. Haldar Justin P.. Low-Rank Modeling of Local k-Space Neighborhoods (LORAKS) for Constrained MRI. IEEE Transactions on Medical Imaging. 2014;33(3):668–681.

28. Shin Peter J., Larson Peder E Z.., Ohliger Michael A., et al. Calibrationless parallel imaging reconstruction based on structured low-rank matrix completion. Magnetic Resonance in Medicine. 2014;72(4):959–970.

29. Haldar Justin P., Zhuo Jingwei. P-LORAKS: Low-rank modeling of local k-space neighborhoods with parallel imaging data. Magnetic Resonance in Medicine. 2016;75(4):1499–1514.

30. Lee Dongwook, Jin Kyong Hwan, Kim Eung Yeop, Park Sung-Hong, Ye Jong Chul. Acceleration of MR parameter mapping using annihilating filter-based low rank hankel matrix (ALOHA). Magnetic Resonance in Medicine. 2016;76(6):1848–1864.

31. Candés Emmanuel J., Sing-Long Carlos A., Trzasko Joshua D.. Unbiased Risk Estimates for Singular Value Thresholding and Spectral Estimators. IEEE Transactions on Signal Processing. 2013;61(19):4643–4657.

32. Veraart Jelle, Novikov Dmitry S., Christiaens Daan, Ades-aron Benjamin, Sijbers Jan, Fieremans Els. Denoising of diffusion MRI using random matrix theory. NeuroImage. 2016;142:394–406.

33. Veraart Jelle, Novikov Dmitry S., Christiaens Daan, Ades-aron Benjamin, Sijbers Jan, Fieremans Els. Denoising of diffusion MRI using random matrix theory. NeuroImage. 2016;142:394–406.

34. Haase A, Frahm J, Matthaei D, Hanicke W, Merboldt K. D. FLASH imaging. Rapid NMR imaging using low flip-angle pulses. Journal of Magnetic Resonance (1969). 1986;67(2):258–266.

35. Dai Weiying, Garcia Dairon, Bazelaire Cedric, Alsop David C.. Continuous flow-driven inversion for arterial spin labeling using pulsed radio frequency and gradient fields. Magnetic Resonance in Medicine. 2008;60(6):1488–1497.

36. Zhao Moss Y., Mezue Melvin, Segerdahl Andrew R., et al. A systematic study of the sensitivity of partial volume correction methods for the quantification of perfusion from pseudo-continuous arterial spin labeling MRI. NeuroImage. 2017;162:384–397.

37. Chang Yulin V., Vidorreta Marta, Wang Ze, Detre John A.. 3D-accelerated, stack-of-spirals acquisitions and reconstruction of arterial spin labeling MRI. Magnetic Resonance in Medicine. 2017;78(4):1405–1419.

38. Carr James C., Simonetti Orlando, Bundy Jeff, Li Debiao, Pereles Scott, Finn J. Paul. Cine MR Angiography of the Heart with Segmented True Fast Imaging with Steady-State Precession. Radiology. 2001;219(3):828–834.

39. Uecker Martin, Lai Peng, Murphy Mark J., et al. ESPIRiT—an eigenvalue approach to autocalibrating parallel MRI: Where SENSE meets GRAPPA. Magnetic Resonance in Medicine. 2014;71(3):990–1001.

40. Lustig Michael, Donoho David, Pauly John M.. Sparse MRI: The application of compressed sensing for rapid MR imaging. Magnetic Resonance in Medicine. 2007;58(6):1182–1195.

41. Blumenthal Moritz, Heide Martin, Holme Christian, et al. mrirecon/bart: version 0.9.00. 2023.

42. Uecker Martin, Ong Frank, Tamir Jonathan I., et al. Berkeley advanced reconstruction toolbox. In: Proc. Intl. Soc. Mag. Reson. Med., International Society for Magnetic Resonance in Medicine; 2015:9; Toronto, ON, Canada.

43. Jenkinson Mark, Bannister Peter, Brady Michael, Smith Stephen. Improved Optimization for the Robust and Accurate Linear Registration and Motion Correction of Brain Images. NeuroImage. 2002;17(2):825–841.

44. Andrews-Hanna Jessica R., Snyder Abraham Z., Vincent Justin L., et al. Disruption of Large-Scale Brain Systems in Advanced Aging. Neuron. 2007;56(5):924–935.

45. Van Dijk Koene R.A., Hedden Trey, Venkataraman Archana, Evans Karleyton C., Lazar Sara W., Buckner Randy L.. Intrinsic Functional Connectivity As a Tool For Human Connectomics: Theory, Properties, and Optimization. Journal of Neurophysiology. 2010;103(1):297– 321.

46. Fonov VS, Evans AC, McKinstry RC, Almli CR, Collins DL. Unbiased nonlinear average age-appropriate brain templates from birth to adulthood. NeuroImage. 2009;47:S102.

47. Fonov Vladimir, Evans Alan C., Botteron Kelly, Almli C. Robert, McKinstry Robert C., Collins D. Louis. Unbiased average age-appropriate atlases for pediatric studies. NeuroImage. 2011;54(1):313–327.

48. Suzuki Yuriko, Clement Patricia, Dai Weiying, et al. ASL lexicon and reporting recommendations: A consensus report from the ISMRM Open Science Initiative for Perfusion Imaging (OSIPI). Magnetic Resonance in Medicine. 2024;91(5):1743–1760.

49. Avants B. B., Epstein C. L., Grossman M., Gee J. C.. Symmetric diffeomorphic image registration with crosscorrelation: Evaluating automated labeling of elderly and neurodegenerative brain. Medical Image Analysis. 2008;12(1):26–41.

50. Tustison Nicholas J., Cook Philip A., Holbrook Andrew J., et al. The ANTsX ecosystem for quantitative biological and medical imaging. Scientific Reports. 2021;11(1):9068.

51. Elliott Maxwell L., Nielsen Jared A., Hanford Lindsay C., et al. Precision brain morphometry using cluster scanning. Imaging Neuroscience. 2024;2:imag–2–00175.

52. Pfaffenrot Viktor, Norris David. Validating NORDIC denoising on high-resolution fMRI data at 7 T. In: Proceedings of 2023 ISMRM & ISMRT Annual Meeting, International Society for Magnetic Resonance in Medicine; 2023:2719; Toronto, ON, Canada.

53. Olesen Jonas L., Ianus Andrada, Østergaard Leif, Shemesh Noam, Jespersen Sune N.. Tensor denoising of multidimensional MRI data. Magnetic Resonance in Medicine. 2023;89(3):1160–1172.

54. Faes Lonike K., Lage-Castellanos Agustin, Valente Giancarlo, et al. Evaluating the effect of denoising submillimeter auditory fMRI data with NORDIC. Imaging Neuroscience. 2024;2:imag–2–00270.

